# Fluence rate-dependent kinetics of light-triggered liposomal doxorubicin assessed by quantitative fluorescence-based endoscopic probe

**DOI:** 10.1101/2024.12.29.630668

**Authors:** Daniel J. Rohrbach, Kevin A. Carter, Dandan Luo, Shuai Shao, Semra Aygun-Sunar, Jonathan F. Lovell, Ulas Sunar

**Affiliations:** Agilent Technologies, Winooski, Vermont, VT 05404, USA; Department of Biomedical Engineering, University at Buffalo, Buffalo, NY 14260, USA; Department of Biomedical, Engineering, Stony Brook University, Stony Brook, NY, 11794, USA

**Keywords:** doxorubicin drug concentration, diffuse reflectance spectroscopy, diffuse fluorescence spectroscopy, light-triggered release, porphyrins, photodynamic therapy, endoscopic probe

## Abstract

Liposomal doxorubicin (Dox), a treatment option for recurrent ovarian cancer, often suffers from suboptimal biodistribution and efficacy, which might be addressed with precision drug delivery systems. Here, we introduce a catheter-based endoscopic probe designed for multispectral, quantitative monitoring of light-triggered drug release. This tool utilizes red-light photosensitive porphyrin-phospholipid (PoP), which is encapsulated in liposome bilayers to enhance targeted drug delivery. By integrating diffuse reflectance and fluorescence spectroscopy, our approach not only corrects the effects of tissue optical properties but also ensures accurate drug delivery to deep-seated tumors. Preliminary results validate the probe effectiveness in controlled settings, highlighting its potential for future clinical adaptation. This study sets the stage for in vivo applications, enabling the exploration of next-generation treatment paradigms for the management of cancer by optimizing chemotherapy administration with precision and control.

## 1. Introduction

Liposomal doxorubicin (Dox) is a commonly utilized therapeutic agent for recurrent ovarian carcinoma, yet its efficacy is often hindered by inefficient drug delivery to tumor cells [1–10]. To address these limitations, nanocarriers such as liposomes have been developed to enhance drug distribution and efficacy. Among these, porphyrin-phospholipid (PoP) liposomes present a unique advantage, allowing controlled drug release triggered by near-infrared (NIR) light with precise spatial and temporal resolution [1,5–8,10–16].

This light-triggered release mechanism not only improves targeted drug delivery but also minimizes systemic side effects by confining the release to specific sites and times. For emerging nanocarriers like PoP-liposomes, obtaining detailed information on local drug concentration distribution is crucial for assessing therapeutic efficacy [5,17–27,27–36]. Both Dox and PoP exhibit fluorescence, which enables the use of fluorescence spectroscopy or imaging for this purpose. However, in turbid media, background optical properties can distort the raw fluorescence signal, making it challenging to accurately quantify drug concentration [24,28,37–46]. Additionally, these optical properties can attenuate the treatment light, delaying drug release.

To overcome the effect of background optical properties, quantitative fluorescence techniques are needed. Several spectroscopic methods have been implemented for quantification of drug concentration *in vivo* [24,28,45,46,37–44]. In this study, we utilized a combination of diffuse reflectance and fluorescence spectroscopy to quantify Dox fluorescence while correcting for variations caused by absorption and scattering at both excitation and emission wavelengths. Using a catheter-based endoscopic probe, we measured the localized concentration of Dox released by NIR light in vitro and in an animal carcass. The probe, which fits into an endoscope’s working channel, provides multispectral data for Dox (∼590 nm) and PoP (∼680–730 nm). This approach enables precise quantification of each component in multi-functional constructs, offering significant potential for intratumoral applications in thick and deep-seated tumors.

## 2. Materials and Methods

### 2.1. Drug Preparation

The details of our PoP Dox (PoP-D) liposome formulation has been described elsewhere [1,12,47,48] . PoP-liposomes incorporated PoP that was in a manner as recently reported [49]. Briefly, PoP-liposomes were synthesized from pyro-lipid through esterification of pyro with lyso-C16-PC, using 1-Ethyl-3-(3-dimethylaminopropyl)carbodiimide (EDC) and 4-dimethylaminopyridine (DMAP) in chloroform. The liposomes were formed by dispersing porphyrin-lipid, PEGylated-lipid, cholesterol, and distearoylphosphatidylcholine in chloroform, followed by solvent evaporation. A 20 mg/mL lipid solution was extruded through a high-pressure lipid extruder with a 250 mM ammonium sulfate solution using polycarbonate membranes of 0.2, 0.1, and 0.08 μm pore size, sequentially stacked and passed through the extruder 10 times. Free ammonium sulfate was removed by overnight dialysis in a 10% sucrose solution with 10 mM HEPES at pH 7. Dox was loaded by incubating the liposomes at 60 °C for 1 hour, achieving a loading efficacy of over 95% as confirmed by G-75 column tests. The self-assembly status and elution position of PoP-liposomes were tracked using 420 nm excitation and 670 nm emission, while Dox was detected using 480 nm excitation and 590 nm emission in a fluorescence plate reader (TECAN Safire).

### 2.2. Instrumentation

The release kinetics of doxorubicin from the PoP-D liposomes was determined using a custom fluorescence spectroscopy setup. A cuvette was placed in a fiber-coupled 4-way holder (Ocean Optics) with one port connected to a 455nm laser (Thorlabs) and a 90-degree port connected to the sensitive channel of a spectrometer (Ocean Optics). The system was controlled with a custom LabView program. For release, a 660nm LED (Mightex) was placed above and focused into the cuvette. Measurements were acquired before PoPD liposomes were added (PBS only) and after each round of treatment light (every 30 seconds for 30 minutes). Three measurements were acquired at each time point for averaging.

To better mimic the *in vivo* case, a custom diffuse optical spectroscopy system combining diffuse reflectance spectroscopy (DRS) and diffuse fluorescence spectroscopy (DFS) was used as shown in **Figures 1a** and **1b** [31,35,48,50–53]. Briefly, the DRS setup consisted of a tungsten-halogen broadband white light (HL-2000-FHSA, Ocean Optics) as the source and the Master channel of the spectrometer as the detector. The light was directed to the target with one source fiber and the diffusely reflected light was collected with one detector fiber (200 μm dia), with the source detector separation (SD) of 260 μm. For the DFS, a 455 nm laser diode was used as the excitation source and the Slave channel of the spectrometer was used as the detector. A 500 nm long pass filter allowed measuring the fluorescence signal (SD=260 μm) along with a “leakage” signal of the excitation light. The reflected excitation light of the laser diode and the reflected white light from DRS were used to correct the raw DFS signal. All 4 fibers were contained in a narrow 18-gauge needle probe as shown in **Figure 1c**.

**Fig. 1.**
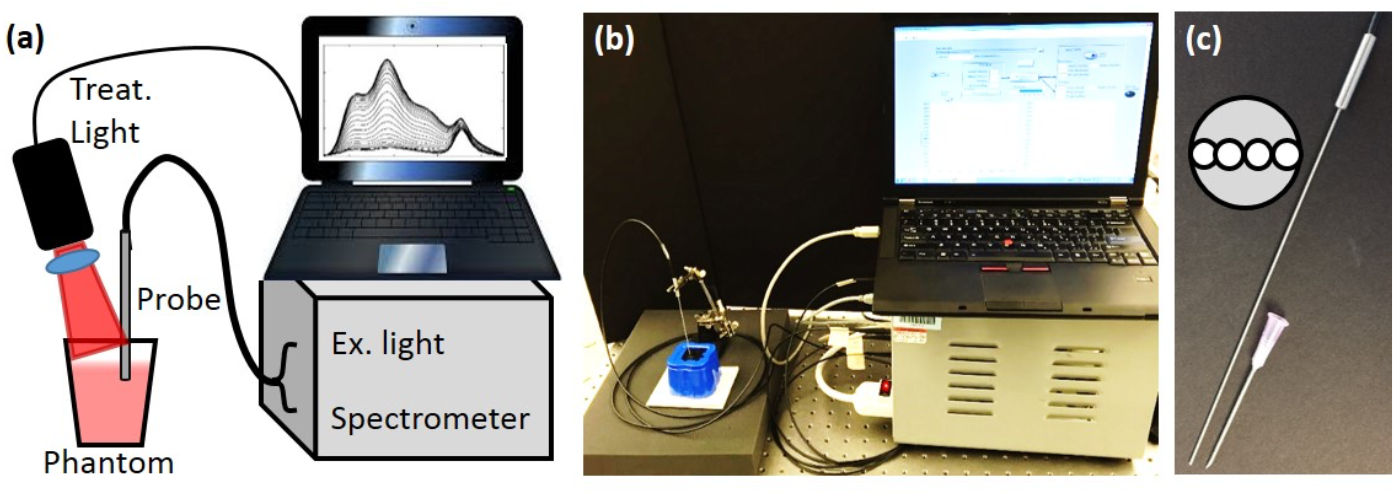
Instrument setup. (a). Diagram of interstitial measurement and treatment system (b) Picture of the setup. (c) The picture of the interstitial needle probe. 18-gauge needle is shown for size comparison. The inset shows the layout of the probe face.

### 2.3 Doxorubicin Quantification

A quantitative fluorescence spectroscopy model was used, similar to Kim *et. al*. [38,41]. Correction is needed to account for the attenuation of the excitation light into the sample as well as the attenuation of emission light out of the sample.

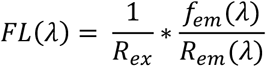

The raw fluorescence signal (*f*_*em*_(*λ*)) was first divided by the white light reflectance spectrum (*R*_*em*_ (λ)), and then scaled by the peak value of the excitation light leakage at 455nm (*R*_*ex*_). The corrected emission (FL(λ)) was then fit to a model of tissue fluorescence, which included background autofluorescence (AF), Dox, and the PoP liposomes (HPPH, also called Pyro),

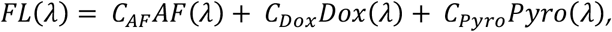

whose corresponding basis spectra are shown in **Figure 2a**. The value *C*_*Dox*_ is then directly related to the true Dox concentration. Sample measurements taken pre- and post-light treatment show the changes in the raw fluorescence due to Dox release (**Fig. 2b**).

**Fig. 2.**
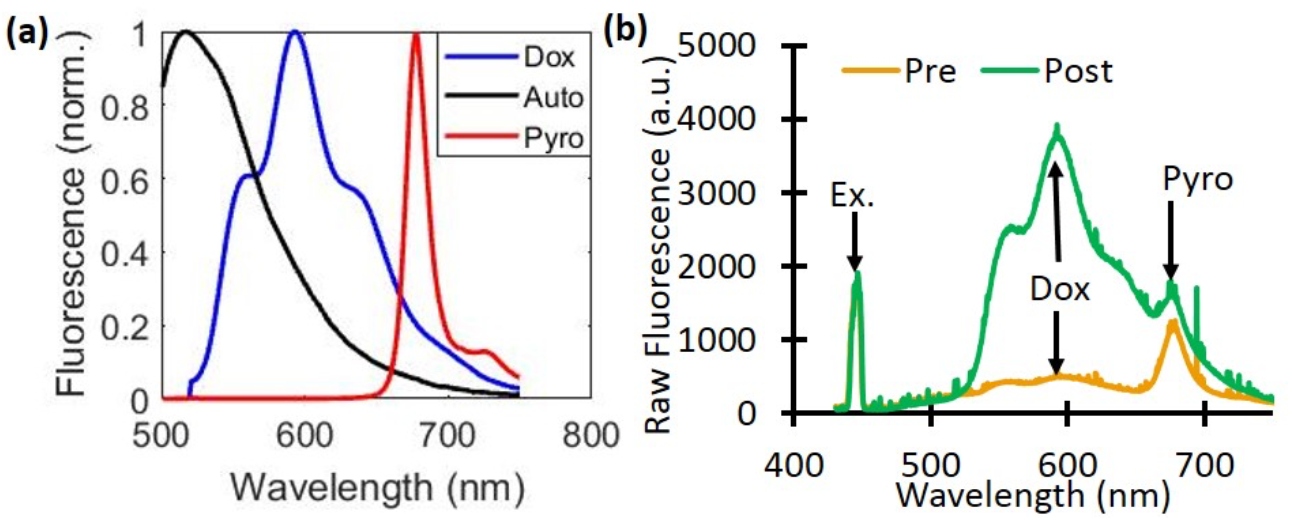
Dox and PoP (Pyro) fluorescence quantification. (a) The basis spectra of autofluorescence (Auto), Doxorubicin (Dox), and PoP were used for quantifying fluorescence concentrations. (b) A representative raw fluorescence signal measured pre- and post-Dox release.

### 2.3 Phantom and animal preparation

Phantoms were prepared using Intralipid 20% (Fresenius Kabi) for scattering and India Ink (Higgins) for absorption. Four phantoms were prepared with different combinations of optical properties: μ_a_ = 0.5 and 1.5 cm^−1^, μ_s_’ = 20 and 30 cm^−1^ at 455 nm; μ_a_ = 0.4 and 1.2 cm^−1^, μ_s_’ = 15.8 and 23.8 cm^−1^ at 590 nm. Each phantom had a total volume of 120 mL. For the calibration of Dox concentration, a stock solution of 0.5 mg/mL free-Dox was used. After baseline reflectance and fluorescence measurements of each phantom, increasing volumes of free-Dox were added to each (400, 800, 1200, 1600, and 2000 μL), with reflectance and fluorescence measurements acquired at each addition.

Dox release was performed in phantoms prepared as above with μ_s_’ = 20 cm^−1^ and μ_a_ = 0.5 and 1.0 cm^−1^ at 455 nm and a total volume of 70 mL. A stock solution of 2.59 mg/mL PoP liposomes was prepared, as described in the previous section. The phantom was placed on a stir plate set to 15 rpm. Simulated interstitial measurements were acquired by placing the needle probe into the liquid phantom to acquire measurements. Fluorescence measurements were acquired before and after the addition of PoP liposomes (200 μL). The stir plate was increased to 30 rpm to ensure a uniform release, and the treatment light (1.72 cm diameter spot) was turned on. Fluorescence measurements were acquired every 4 minutes.

Next, a recently sacrificed nude mouse was used for mimicking *in vivo* measurements to check and optimize the signal. The mouse was injected subcutaneously with 50 μL of a lightly scattering medium (μ_s_’ = 5 cm^−1^ at 455 nm) containing 22 μg/mL PoP-Liposomes to mimic a tumor under the surface and reflectance and fluorescence spectroscopy measurements were performed. Three measurements were acquired for pre-release baseline levels by placing the probe against the tumor-mimicking sample. The treatment light (1.0 cm diameter) was directed onto the injection site, covering it completely. Fluorescence measurements were acquired every 4 minutes to assess the release kinetics during treatment.

## 3. Results and Discussion

Calibration with free-Dox provided the conversion between corrected fluorescence values and Dox concentrations as shown in **Figure 3. Figure 3a** indicates a representative schematics of the four phantoms and their optical properties. **Figure 3b** shows the mean raw fluorescence signal for all four phantoms at each titration of Dox. The large error bars represents the standard deviation of the four phantoms and highlight the effects of background optical properties on the raw fluorescence. **Figure 3c** shows the corrected fluorescence (*C*_*Dox*_ from Equation 2). The corrected fluorescence has much smaller error bars, highlighting the improved quantification. In addition, after the correction, the slope of the line directly relates the relationship between the corrected Dox fluorescence to the true Dox concentration, so that one can extract the local Dox concentration for unknown sample.

**Fig. 3.**
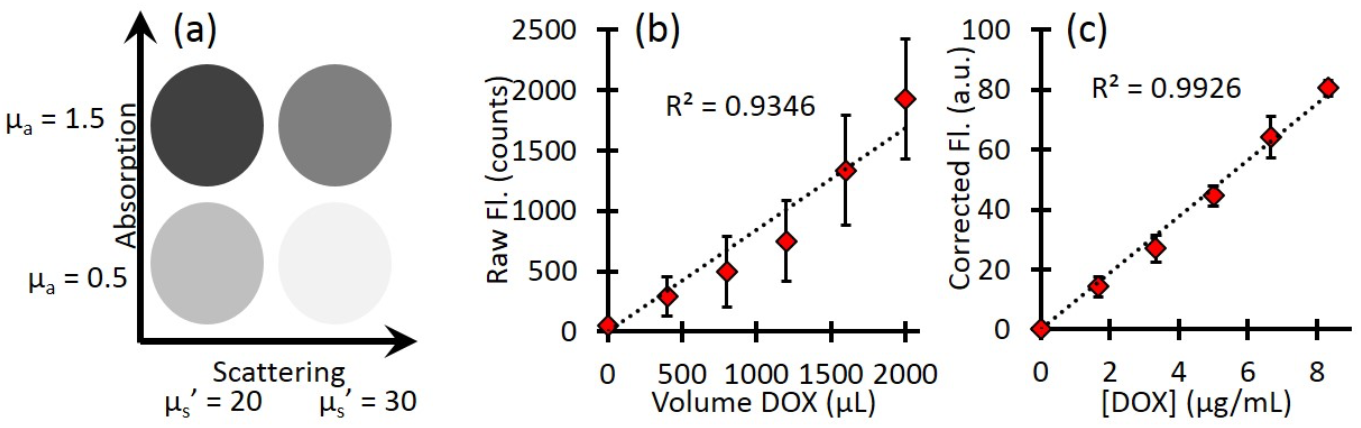
Phantom calibration. (a) A schematic of the phantom optical properties (b) Raw Dox fluorescence counts show the effect of background optical property variation. Error bars represent the standard deviation between phantoms. (c) Corrected fluorescence shows a linear response to increasing Dox concentration and much less variation between phantoms. Error bars represent the standard deviation between phantoms.

Next, we investigate the release kinetics with respect to light irradiation fluence rate. Our fluorescence spectroscopy measurements showed a difference in drug release kinetics with fluence-rate dependent manner (**Fig. 4a**). The lower fluence rate only started releasing Dox after 450 seconds and was fully released by 900 seconds while the higher fluence rate started releasing Dox after 270 seconds and was fully released by 510 seconds. This is a difference in release rate of 450 seconds for the low fluence rate and 240 seconds for the high fluence rate. However, when plotted as a function of light dose (mW/cm^2^) the two curve are much more similar (**Fig. 4b**). As **Figures 4c** and **4d** indicate, there was a decrease in the Pyro fluorescence as well, approximately from 4000 counts to 3000 counts, indicates the photobleaching of the porphyrin component.

**Fig. 4.**
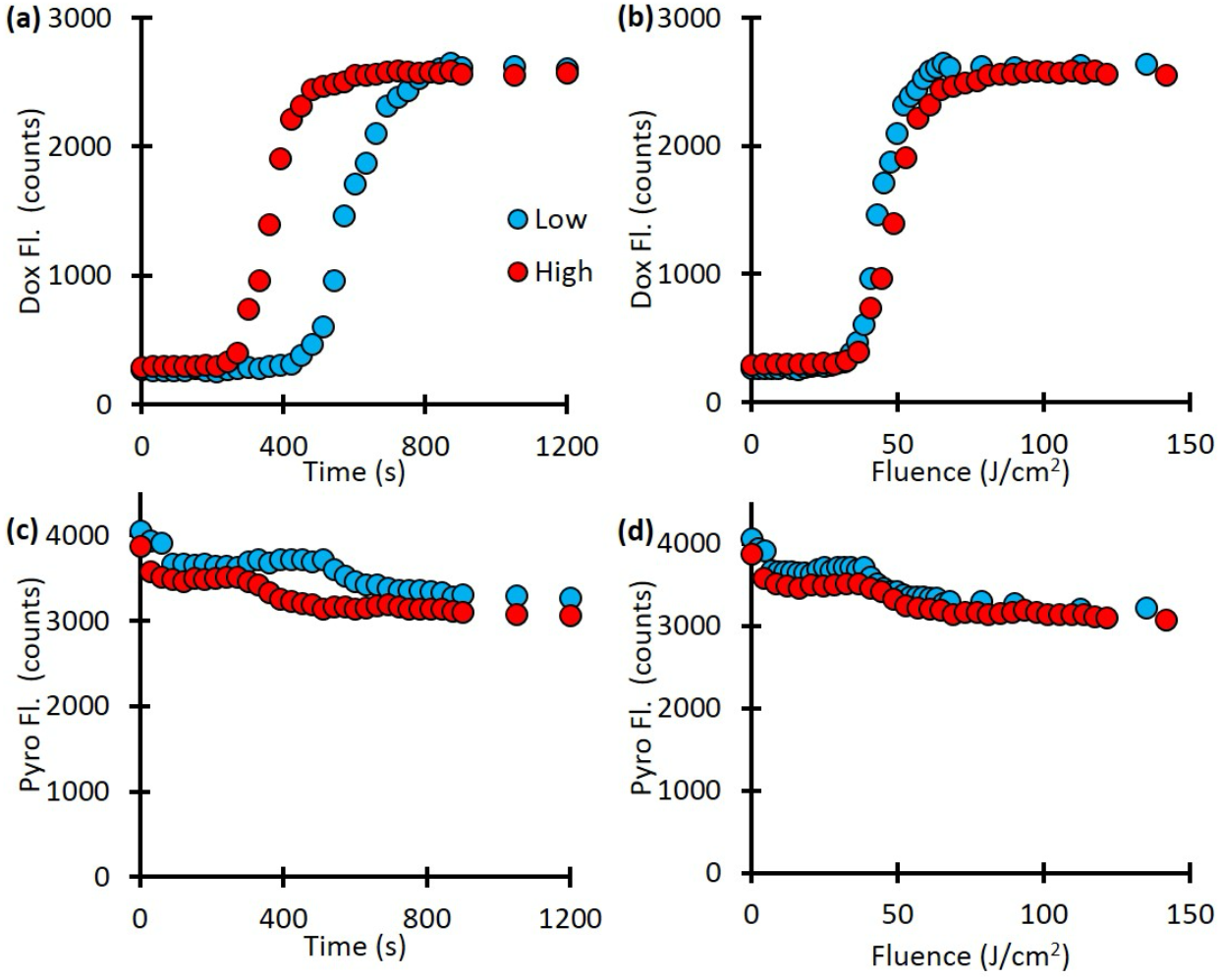
The release kinetics. The time to complete release depends on the fluence rate. (a) Dow release as a function of time for different fluence rates. (b) Dox release as a function of fluence rate. (c) Pyro-fluorescence signal as a function of time. (d) Pyro-fluorescence signal, as a function of fluence rate, indicates the photobleaching of the porphyrin component.

The fluence rate can be precisely controlled for optically clear phantoms in a cuvette. However, for turbid media, the true local-site fluence is unknown since the background optical properties will affect the delivered treatment light. The background optical properties will also affect the determination of fluorophore concentration. To mimic this case, we check the kinetics for the high and low-absorption cases. As **Figure 5a** shows, the raw fluorescence for phantoms with the same PoPD liposome concentration showed different rates of release as well as different values at full release. The phantom with lower background absorption showed a final Dox signal of 2007.1 ± 44.8 counts, while the higher absorbing phantom showed 1656.5 ± 46.1 counts, a difference of 17.5%. However, after quantifying the Dox concentration (**Fig. 5b**), the full release for the low and high-absorbing phantoms was 11.4 ± 1.5 μg/mL vs. 11.5 ± 1.1 μg/mL, respectively, a difference of less than 1%. As **Figure 5c** shows, the signal from Pyro (PoP, HPPH) decreased slightly throughout treatment for both the low and high-absorbing phantom (26.8% and 10.4%, respectively). This decrease is likely due to the photobleaching of the Pyro.

**Fig. 5.**
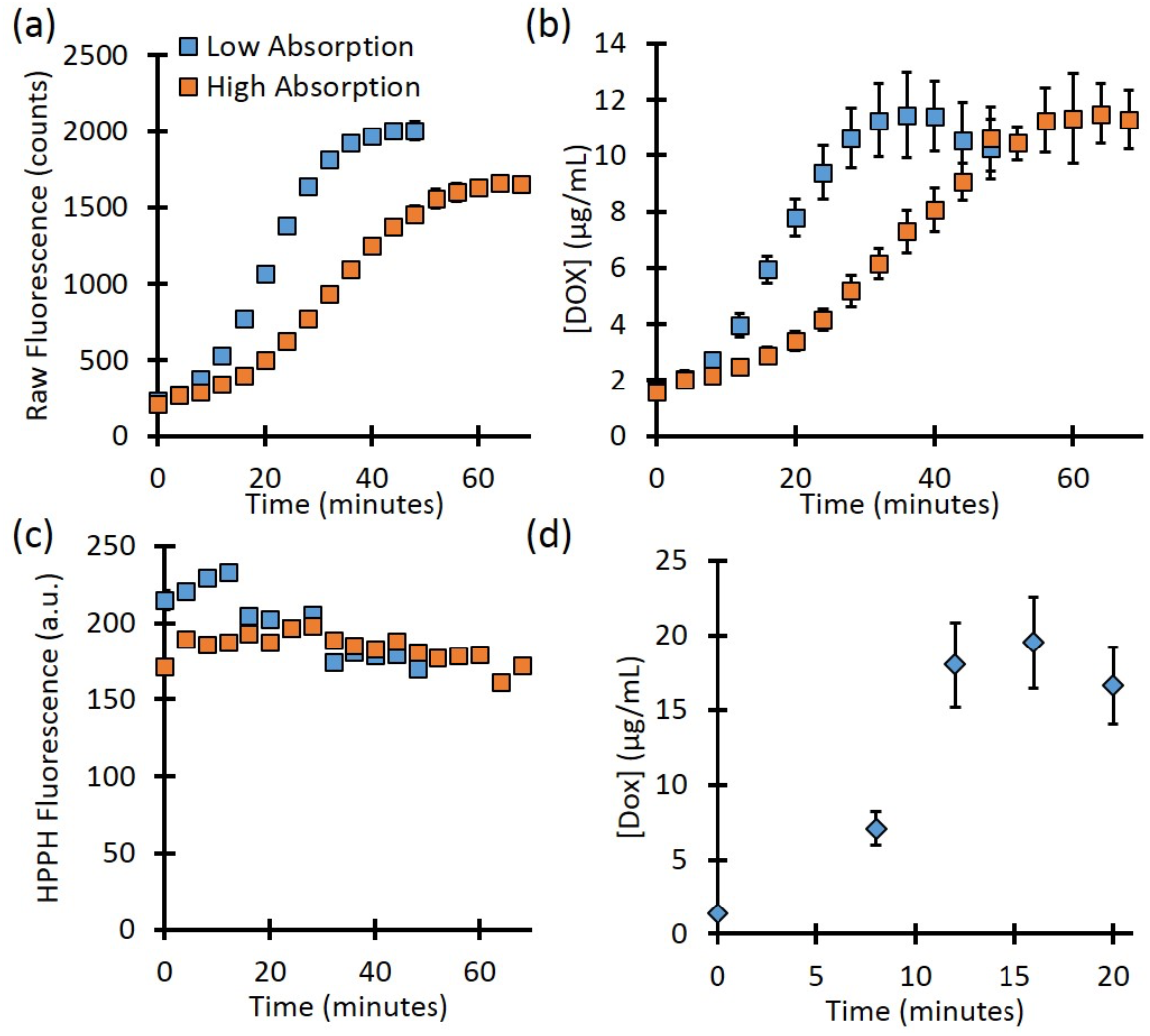
(a) Background absorption differences showed different final values of raw fluorescence after release. (b) Corrected fluorescence fitting showed the same final dox concentration after release for the two phantoms. (c) Pyro (PoP, HPPH) fluorescence showed a slight decrease throughout treatment. (d) The release in a mouse model.

## 3. Discussion

For the mouse experiment, the expected value of Dox concentration was 22 μg/mL based on the concentration of the preparation. The corrected fluorescence fitting determined the Dox concentration to be 19.5 μg/mL, a difference of 11.4%. This difference could be due to the partial volume effect due to the skin layer. It is clear that Dox accumulates at the local site with time, which is the main aim for an effective cell kill. It is interesting to note that the PoP amount was photobleached with time. This aligned well with the previous observations of others and ours related to photodynamic therapy (PDT) treatment of tumors with HPPH. Since the release light intensity is on the same order of magnitude as PDT, where photosensitizers (like HPPH) are photobleached due to porphyrin, oxygen, and light interaction. It would be interesting to investigate the combination effect (PDT and Dox-chemo) for improved cell-killing efficacy.

An important part of the fluorescence quantification is the correction for attenuation of excitation light. This was accomplished by allowing some excitation light to “leak” into the detector fiber. This leakage corresponds to the reflected excitation light and depends on the optical properties at the excitation wavelength. Kim et al. used a similar process for fluorescence quantification, but instead of measuring the ***R***_***ex***_, they calculated ***R***_***ex***_ using knowledge of the tissue optical properties. For our method it is not necessary to measure the tissue optical properties, which enables the fluorescence quantification to be performed very quickly. However, in cases of very high absorption or longer source detector separations where the excitation leakage may not be seen, the calculation method will provide more reasonable estimates of ***R***_***ex***_. In these studies, the ***R***_***ex***_ signal was always seen.

In our previous work, we showed that spatial frequency domain imaging (SFDI) can quantify the release of doxorubicin [1,47,48]. While the SFDI technique is well-suited for superficial surfaces, it may not work for tumors located deeper inside the body. In those cases, quantitative fluorescence spectroscopy with a needle probe can be used.

However, this approach comes with several limitations. While the catheter-based probe provides precise point measurements, it lacks the capability for spatially resolved imaging over larger areas, which could be beneficial for identifying heterogeneous drug distribution in tumors. The system’s ability to measure fluorescence at depths is limited due to limited source-detector separation (here 260 μm). We used the animal carcass, which may not fully replicate the heterogeneity and complexity of in vivo cases. The study focused on a specific set of light parameters (fluence rate and dose). A broader exploration of different fluence rates and dosimetry schemes is needed to establish universally applicable guidelines for light-triggered drug release. To address these limitations, future research could explore multi-modal imaging approaches combining spatial resolution with the point-based probe, validate the methodology in larger and more clinically relevant animal models, and conduct pilot clinical studies to assess feasibility and efficacy in humans.

## 5. Conclusions

In this work, we present results from a custom-built DRS and DFS system for monitoring absolute Dox and PoP concentration changes during light-triggered release in phantoms and in a mouse carcass. We corrected for background optical properties to obtain the true dox concentration. Quantitative Dox concentration release kinetics showed a faster release with respect to laser irradiation. These results show the potential for interstitial measurements to assess Dox release in deeply seated tumors where wide-field imaging techniques cannot reach.

## Acknowledgments

The authors acknowledge the funding support from NIH/NCI R01 (5R01CA243164).

## Author Contributions

U.S. conceived and designed the experiments; D.J.R. performed the experiments; D.J.R. analyzed the data; D.J.R., U.S., SAS, and J.F. Lovell wrote the paper.

## Institutional Review Board Statement

“Not applicable.”

## Informed Consent Statement

“Not applicable.”

## Data Availability Statement

The data that support the findings of this study are available from the corresponding author upon reasonable request.

